# Single Cell Transcriptomic Modelling of the Fallopian Tube Epithelium Identifies Cellular Specialisation, Novel Differentiation Trajectories, and Gene Network Associations with Ectopic Pregnancy

**DOI:** 10.1101/2024.12.20.629653

**Authors:** Lily I Wright, Ivan Wangsaputra, Terence Garner, Megan C Sharps, Roger Sturmey, Peter T Ruane, Adam Stevens

## Abstract

**STUDY QUESTION:** Can network modelling of single cell transcriptomic data identify cellular developmental trajectories of fallopian tube (FT) epithelium and reveal functional and pathological divergence from the endometrium?

**SUMMARY ANSWER:** A bidirectional secretory and ciliated differentiation trajectory was apparent from a novel OVGP1+ progenitor population of FT epithelial cells. A causal network model of whole transcriptome action in the FT and endometrium revealed specific functional divergence between secretory cells of these tissues. The network model reflected the latest ectopic pregnancy genome wide association study (GWAS), invoking *MUC1* and other candidate genes in mature secretory cells for ectopic and eutopic implantation.

**WHAT IS KNOWN ALREADY:** The fallopian tube forms the in vivo peri-conceptual environment, which has a significant impact on programming offspring health. The fallopian tube epithelium establishes this environment, however the epithelial cell types are poorly characterised in health and disease.

**STUDY DESIGN, SIZE, DURATION:** Publicly available benign FT single cell RNA sequencing (scRNA-seq) samples from thirteen women across three studies were combined. Endometrial scRNA-seq samples from thirteen women from one study were used to demonstrate transcriptomic differences between the epithelia of the two tissues. Network models of transcriptomic action were constructed with hypergraphs.

**PARTICIPANTS/MATERIALS, SETTING, METHODS:** A meta-analysis of FT scRNA-seq samples was performed to identify epithelial populations. Differential gene expression assessed differences between fallopian tube and endometrial epithelial scRNA-seq data. Functional differences between secretory cells in the tissues were characterised using hypergraph models. To identify associations with ectopic pregnancy, expression quantitative trait loci (eQTLs) from a recent GWAS were mapped onto the network models.

**MAIN RESULTS AND THE ROLE OF CHANCE:** Epithelial cells (n=14,360) were clustered into 8 secretory and ciliated epithelial populations in the meta-analysis of 3 scRNA-seq datasets. A novel *OVGP1*+ epithelial progenitor cell was also identified, and its bi-directional differentiation to mature secretory or mature ciliated populations was mapped by RNA velocity analysis. This progenitor exhibited a high velocity magnitude (12.47) and low confidence (0.69), a combination strongly indicative of multipotent progenitor status. Comparing FT epithelial cells with endometrial epithelial cells revealed 5.3-fold fewer shared genes between FT and endometrial glandular secretory cells than between FT and endometrial ciliated cells, suggesting functional divergence of secretory cells along the reproductive tract.

Hypergraphs were used to identify highly coordinated regions of the transcriptome robustly associated with functional gene networks. In the FT secretory cells, these networks were enriched for lipid (FDR<0.002) and immune (FDR<0.00007) related pathways. We mapped eQTLs from a GWAS meta-analysis of 7070 women with ectopic pregnancy over a range of significance (P = 1.68 x 10^-21^– 5.8 x 10^-4^) to the hypergraphs of FT and endometrium. Of the 22 genes present in the hypergraphs, 13 of these clustered as highly coordinated genes. This demonstrated the functional importance of *MUC1* in the FT and endometrium, (GWAS Study P = 5.32×10^-9^) and identified additional genes (*SLC7A2, CLDN1, GLS, PEX6, PLXNA4, NR2F1, CLGN, PGGHG, ANKRD36*) implicated in ectopic pregnancy and eutopic pregnancy.

**LIMITATIONS, REASONS FOR CAUTION:** The sample size of reproductive age women was limited in previous studies, and though causal network modelling was used and previous mechanistic data supports candidate gene involvement, no in vitro or in vivo validation of candidate was performed.

**WIDER IMPLICATIONS OF THE FINDINGS:** These findings consolidate the existing single cell transcriptomic datasets of the FT to provide a comprehensive understanding of epithelial populations and define functionally distinct secretory cells that contribute to the peri-conceptual environment of the FT. We further implicate the role of MUC1 and secretory cells in ectopic pregnancy and suggest future targets for investigating embryo implantation in the FT and endometrium.

## Introduction

The fallopian tube (FT) is the site of early reproductive processes in vivo, facilitating gamete transport, fertilisation and pre-implantation embryo development. The peri-conceptual environment provided by the FT has a critical hand in programming offspring health, as underscored by metabolic deficiency in animal studies and assisted reproductive treatment outcomes in human[1, 2] Moreover, FT dysfunction can cause infertility and ectopic pregnancy[3, 4]. Despite this fundamental role in reproductive health and developmental programming, the biological mechanisms underpinning FT function in health and disease remain underexplored [5, 6].

The FT is divided into four regions; distally, the fimbriae, infundibulum and ampulla, and proximal to the uterus, the isthmus. It is composed of a thin outer smooth muscle layer, the myosaplinx, with the endosalpinx consisting of stroma and a simple columnar epithelium of secretory and multi-ciliated cells lining the lumen. The myosalpinx and ciliated epithelium combine to move tubal fluid along the tract and the secretory cells of the epithelium produce secretions which provide the peri-conceptual environment for gametes and pre-implantation embryos [7, 8]. FT fluid has been studied ex vivo and in vivo, demonstrating its effect on embryo metabolism and DNA methylation [9–11]. Moreover, recent in vivo work in mice confirms the role of ciliated cells in moving fluid thus contributing to gamete/embryo transport [12, 13].

To highlight the FT as a distinct functional tissue, we compare FT and endometrial epithelial cells in the secretory phase. The FT and uterus share common embryological origins and exist as a continuum, despite their anatomical proximity they have significantly different functions in reproduction[14, 15]. Unlike the endometrium, the FT lumen does not undergo a receptivity period, yet it is susceptible to embryo implantation in 1-2% of pregnancies [16]. Tubal ectopic pregnancy is a common but serious condition, though subsequent placentation is not viable; it can cause severe haemorrhage when undetected.

Several single cell RNA sequencing (scRNA-seq) datasets investigating ovarian cancer, hydrosalpinx and ectopic pregnancy have characterised cell types of the FT epithelium [17–19]. These datasets highlight alternate hypotheses of epithelial differentiation in the FT. Dinh et al. [19] modelled two differentiation pathways, where secretory progenitor cells give rise to a mature secretory population, and an alternate ‘epithelial to mesenchymal transformation’ cell type gives rise to both ciliated epithelial cells and stromal cells. Conversely, Ulrich et al. [18] describe secretory cells differentiating to ciliated cells via a highly proliferative transitioning population.

Single-cell transcriptome datasets also enable the analysis of developmental trajectories and causal modelling, allowing us to infer functional and pathological gene networks within cell populations of the FT. By constructing hypergraphs from gene expression matrices, we construct gene network models which capture relationships among groups of genes rather than pairs of genes [20]. Identifying these higher order gene interactions, allows us to determine genes with a high number of connections to the rest of the transcriptome, suggesting their functional biological significance [21]. Hypergraph models have been shown to capture biological relevance and causality in omics datasets of embryo implantation[22], rare disease[23], and developmental programming [24].

Here we perform a meta-analysis of healthy FT scRNA-seq data from 13 women to examine FT epithelial cell sub-populations and differentiation in greater depth and identify clinically relevant functional differences between FT and endometrium in the context of tubal ectopic pregnancy.

## Methods

### scRNA-seq data and samples

Published FT scRNA-seq datasets were obtained from GSE183837 [17], GSE151214[19] and GSE178101 [18]. Published endometrial scRNA-seq dataset was obtained from E-MTAB-10287 [25]. Cadaveric samples and those from patients with diagnosed malignancies were excluded. FT data from 13 patient samples, and endometrial data from 13 patient samples from were included in final data analysis. Publicly available demographics are shown in Supplementary Table 1.

### Raw Data Processing

Raw data processing was performed using a standardised methodology. FASTQ files were processed using Cell Ranger (7.1.0) analysis pipeline and aligned to GRCh38 reference genome to generate single cell gene expression count files (51). For GSE183837 [17], FASTQ files were processed using HISAT2 (2.2.1) and SAMtools (1.9). Velocyto (0.17) was used to generate all unspliced and spliced count matrices for RNA velocity analysis [26].

### scRNA-seq analysis

scRNA-seq data analyses were performed using single cell gene expression count files in Python (3.11.4). Scanpy package (1.9.3) was used for all pre-processing, quality control, dimensionality reduction, visualisation and marker gene identification steps [27].

### Filtering, preprocessing and dimensionality reduction

For all data analysis, cells with <200 genes expressed, and genes expressed in <3 cells were removed, following standard parameters of the Scanpy package [27]. Cells with >850,000 total gene counts, >30% mitochondrial gene counts, and >8000 genes in the count matrix were also removed. To batch correct and integrate multiple datasets, datasets were concatenated, highly variable genes were selected with a batch factor of patient, ComBat [28] batch correction was used with dataset as a batch factor.

For individual FT datasets dimensionality reduction and clustering, nearest neighbour graphs were constructed using 8 principal components and 10 neighbours. Leiden clustering was performed between 0.4-0.5 resolution. To produce the combined FT cellular atlas, raw count files of individual FT datasets were concatenated and then filtered and pre-processed. Nearest neighbours graph was constructed using 9 principal components, 10 neighbours, Leiden clustering was run at 1.7 resolution. FT epithelium was sub-clustered by cropping data to only Ciliated and Secretory Epithelial Cells. Dimensionality reduction and clustering were re-computed, and nearest neighbours graph was constructed using 7 principal components and 90 neighbours.

To resolve endometrial cell clusters, nearest neighbours graph was constructed with 15 principal components and 300 neighbours. Epithelium was sub-clustered by cropping data to only Sox9+ Epithelial, Lumenal Epithelial, Glandular Epithelial and Ciliated cells. Dimensionality reduction and clustering were re-computed, nearest neighbours graph was constructed with 25 principal components and 200 neighbours. Leiden clustering was run at resolution 1.

To compare endometrial and FT epithelial cells, epithelial datasets were concatenated, batch corrected, and dimensionality reduction was re-computed. Nearest neighbours graph was constructed with 25 principal components and 200 neighbours. The endometrial and FT full cellular atlas was constructed by concatenating raw datasets, batch corrected, dimensionality reduction and clustering were computed to identify cell types. Nearest neighbours graph was constructed with 25 principal components and 250 neighbours. Leiden clustering was run at resolution 2.

In the FT, major cell types and epithelial sub-populations were annotated by inspection of literature-based marker genes, and cluster defined marker genes. In the endometrium, cell types were annotated using the marker genes reported by Garcia-Alonso et al [25]. Differentially expressed marker genes were identified for each cluster using Wilcoxon test. ScVelo package (0.2.5) was used for RNA velocity analysis plotting functions and computing velocity magnitude and confidence [29]. Proportional Venn diagrams were generated using BioVenn [30].

### Statistical analysis

Linear regression was used to compare proportions of epithelial sub-populations between menstrual cycle phases in FT and endometrial epithelium, in GraphPad Prism (9.5.1). Differential gene expression was tested using Wilcoxon rank sum in Scanpy (1.9.3)

### Hypergraph analysis

Hypergraphs were generated to investigate functional relationships between the differentially expressed genes between the FT and endometrial epithelia, and the rest of the transcriptome in the FT, glandular and luminal endometrium. All hypergraph analyses were performed in R. Correlation matrices were constructed using top 600 differentially expressed genes (DEGs) between secretory phase, secretory epithelial cells of the FT and endometrium as nodes, and the remaining genes from the integrated full cellular atlas of either the FT, glandular or luminal endometrium as edges.

Standard deviation of the correlation matrices were calculated to set a threshold which correlation matrices were binarized against. Matrix multiplications were performed to generate an adjacency matrix where values represent correlations between the defined differentially expressed genes and rest of the transcriptome for each tissue, these were plot as heatmaps where higher counts indicated a higher number of correlations. Hierarchical clustering of the DEGs was performed to identify central clusters of genes with the highest number of correlations, suggesting groups of functionally active genes. Central clusters of functionally active genes were identified for each hypergraph constructed from FT, luminal or glandular epithelium. To compare functionally active genes between the three tissue epithelia, shared and unshared genes of the respective central clusters were identified and proportional Venn diagrams constructed.

### Gene Ontology

Webgestalt 2019 [31] Over Representation Analysis of GeneOntology Biological Process database was used to assess hypergraph central clusters for each tissue. Default parameters were used, False Discovery Rate (FDR) and P values were presented as -log10 using R.

### Tubal Ectopic Pregnancy Genome Wide Association

To identify genes associated with ectopic pregnancy which retain relevance but did not reach the p-value defined by the GWAS study [32], GCST90272883 [33] GWAS summary statistics were downloaded from GWAS catalog. To identify genes where expression is altered, variantIDs of lowest 10,000 p-values were called from GTEx Portal API V2 singleTissueEqtlByLocation, accessed using Python (3.11.4). All human tissues bar nervous system were searched. Summary statistic p-values of the GWAS genes identified in the 600 differentially expressed hypergraph genes were compiled in a violin plot.

To determine significance of ectopic associated GWAS genes across the three tissue hypergraphs, row sums, or degree centrality, of each DEG were calculated. This represents the number of direct connections with the rest of the transcriptome. Row sums were ranked, using an average for split ranks. Ranks of GWAS genes were identified and plot for each tissue.

### Data Availability

Fallopian Tube scRNA-seq datasets are available at GEO under GSE183837 [17], GSE151214[19] and GSE178101 [18]. Endometrial scRNA-seq dataset is available at ArrayExpress under E-MTAB-10287 [25].

## Results

### Meta Analysis of Fallopian Tube Single Cell RNA-seq datasets

Initial clustering of individual fallopian tube (FT) datasets to assess concordance of cell types identified 5 distinct cell populations in the Hu et al.[17] dataset, 10 in the Dinh et al. [19] dataset, and 9 in the Ulrich et al. [18] dataset (Fig 1a,b,c). All datasets contained expected immune populations, secretory and ciliated epithelia, supporting fibroblast, endothelial and smooth muscle cells.

**Figure 1.**
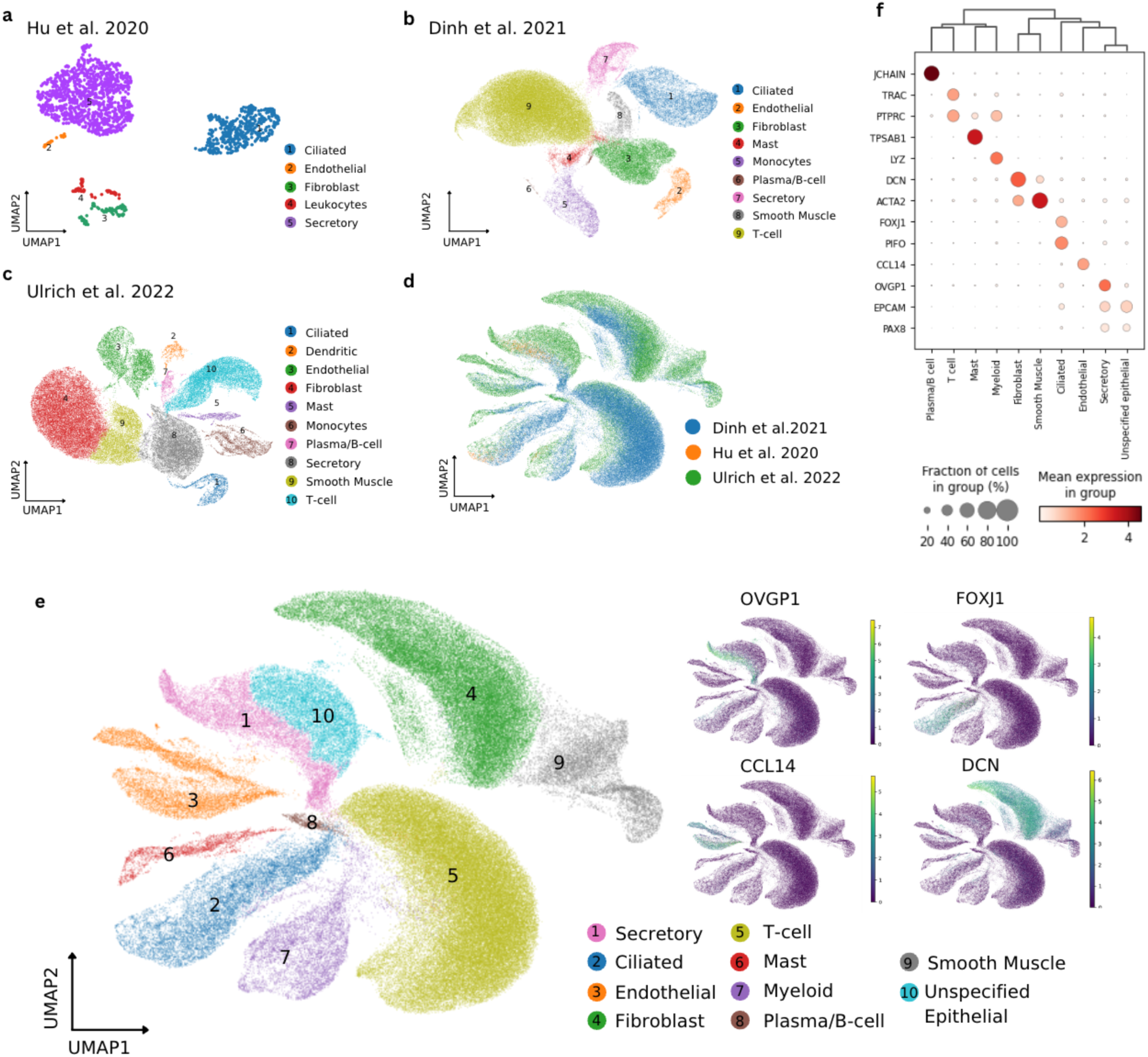
Meta-analysis of Fallopian Tube (FT) scRNA-seq Datasets. **a,** UMAP projection of cell types in Hu et al. scRNA-seq dataset. b, UMAP projection of cell types in Dinh et al. scRNA-seq dataset. c, UMAP projection of cell types in Ulrich et al. scRNA-seq dataset. d, UMAP projection of 3 integrated FT scRNA-seq datasets coloured by Author. e, UMAP projection of 3 integrated FT scRNA-seq datasets, coloured by cell type, cell type marker gene expression is shown, OVGP1 – secretory epithelial cell, FOXJ1-ciliated epithelial cell, CCL14 – endothelial cell, DCN – Fibroblast. f, Dotplot illustrating mean gene expression and fraction of cells expressing marker genes in key cell types of integrated FT cellular atlas.

To perform a meta-analysis of FT scRNA-seq data, the three raw datasets were integrated to a combined matrix of 142,339 cells (Fig 1d,e). Clustering identified 9 major cell types which were annotated according to marker gene expression; 4 immune cell populations, Plasma/B cells ( *JCHAIN+),* T-cells (*TRAC+),* Mast cells (*TPSAB1+),* Myeloid cells *(LYZ+),* Fibroblast (*DCN+),* Endothelial (*CCL14+*), Smooth Muscle (*ACTA2+)*, and Secretory (*OVGP1+*) and Ciliated (*FOXJ1+*) epithelial cells (Fig 1f). A large population of cells marked by classical epithelial markers (E*PCAM, PAX8, KRT18, MMP7*) but were *OVGP1*-negative and so were labelled unspecified epithelial. To understand FT regulation of the peri-conceptual environment, we set out to characterise epithelial cells in more depth.

### The Fallopian Tube Epithelium Contains 8 Populations of Ciliated, Secretory and Progenitor Cells

To focus on epithelial cells, these cells were subset and re-clustered. Initially, twelve clusters were identified among the 14,360 epithelial cells (Fig 2a). Examining top marker genes for each cluster (Fig 2b); Cluster 10 was annotated Mature Secretory-1, marked by high expression of *OVGP1,* a secretory marker, and *LGR5,* a stemness marker (*LGR5+, OVGP1+)*. Clusters 9, 3 and 5 were combined to form a Mature Secretory-2 population (*OVGP1+, LGR5-*). Cluster 11 expressed *OVGP1* and *PIFO*, a marker of primary cilia, but did not express *FOXJ1,* a multi-ciliation marker, so was annotated Mature Secretory (Primary cilia) (*OVGP1+, PIFO+, FOXJ1-, SPAG6+*).

**Figure 2.**
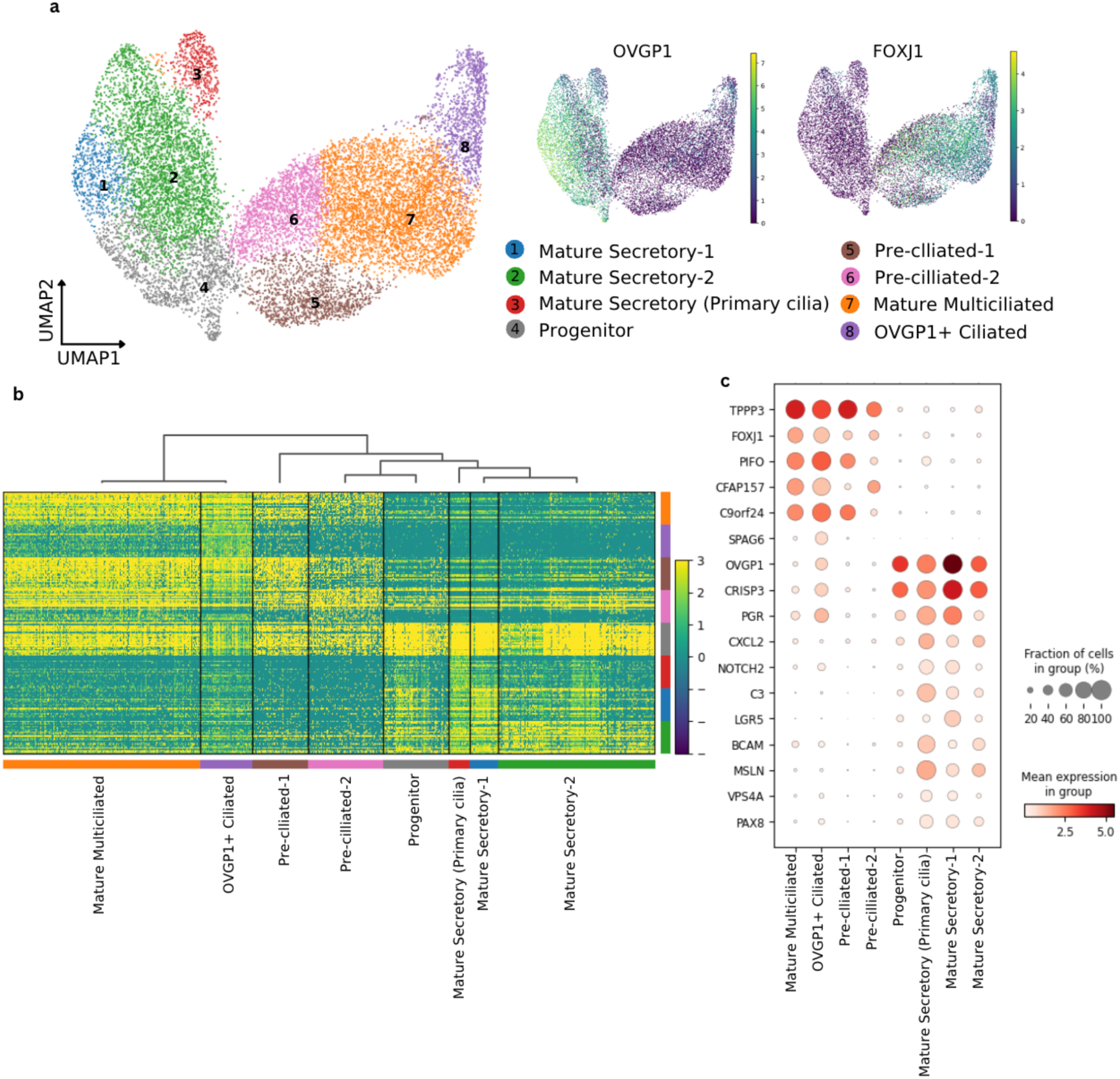
14,630 Fallopian Tube (FT) Epithelial Cells Cluster into 3 Secretory, 1 Progenitor and 4 Ciliated Populations. a, UMAP projection of FT epithelial cells from of Hu et al., Ulrich et al., Dinh et al., datasets. Showing 8 populations, UMAP of OVGP1 expression marking secretory cells, and FOXJ1 expression marking ciliated cells. b, Heatmap of top 25 marker genes for each cell population, genes represented on x-axis and clusters of cell populations on y-axis. c, Dotplot illustrating mean gene expression and fraction of cells expressing key marker genes in populations of FT epithelial cells.

Clusters 0, 4 and 7 were combined to form Mature Multiciliated population (*FOXJ1+, OVGP1-, PIFO+*). Clusters 1 and 6 both expressed lower levels of *FOXJ1* than mature multiciliated clusters, and the absence of some key ciliation genes, such as *CFAP157*, or *C9ORF24*, suggesting they may be developing ciliated cells. Cluster 1 was therefore annotated Pre-ciliated-2 (*PIFO+, CFAP157-, C9orf24+, OVGP1-*) and Cluster 6 was annotated Pre-ciliated-1 (*PIFO+, CFAP157+, C9orf24-, OVGP1-*). Cluster 2 expressed secretory markers, but its proximity to ciliated clusters suggested it may represent a progenitor or transitional population, and was annotated Progenitor (*OVGP1+ C3-, PAX8-, FOXJ1-, PIFO-)*. This allowed us to conclude epithelial cells in the human fallopian tube can be assigned into one of 3 Mature Secretory populations, 4 Ciliated populations or a Progenitor population.

### RNA Velocity Analysis Suggests a Bidirectional Development of Mature Secretory and Ciliated Epithelial Cells in the Fallopian Tube Epithelium

Next, the differentiation trajectory of FT epithelia was assessed. Using RNA velocity analysis[29], a velocity vector is calculated for each cell and the future state of the cell can be inferred from the direction of the velocity vector. By plotting these velocity vectors, it was possible to map a bi-directional developmental trajectory with the progenitor population acting as a source, differentiating into terminal Mature Secretory or Mature Multiciliated populations. The progenitor cells feed into the Pre-Ciliated and then to the Mature Multiciliated populations (Fig 3a).

**Figure 3.**
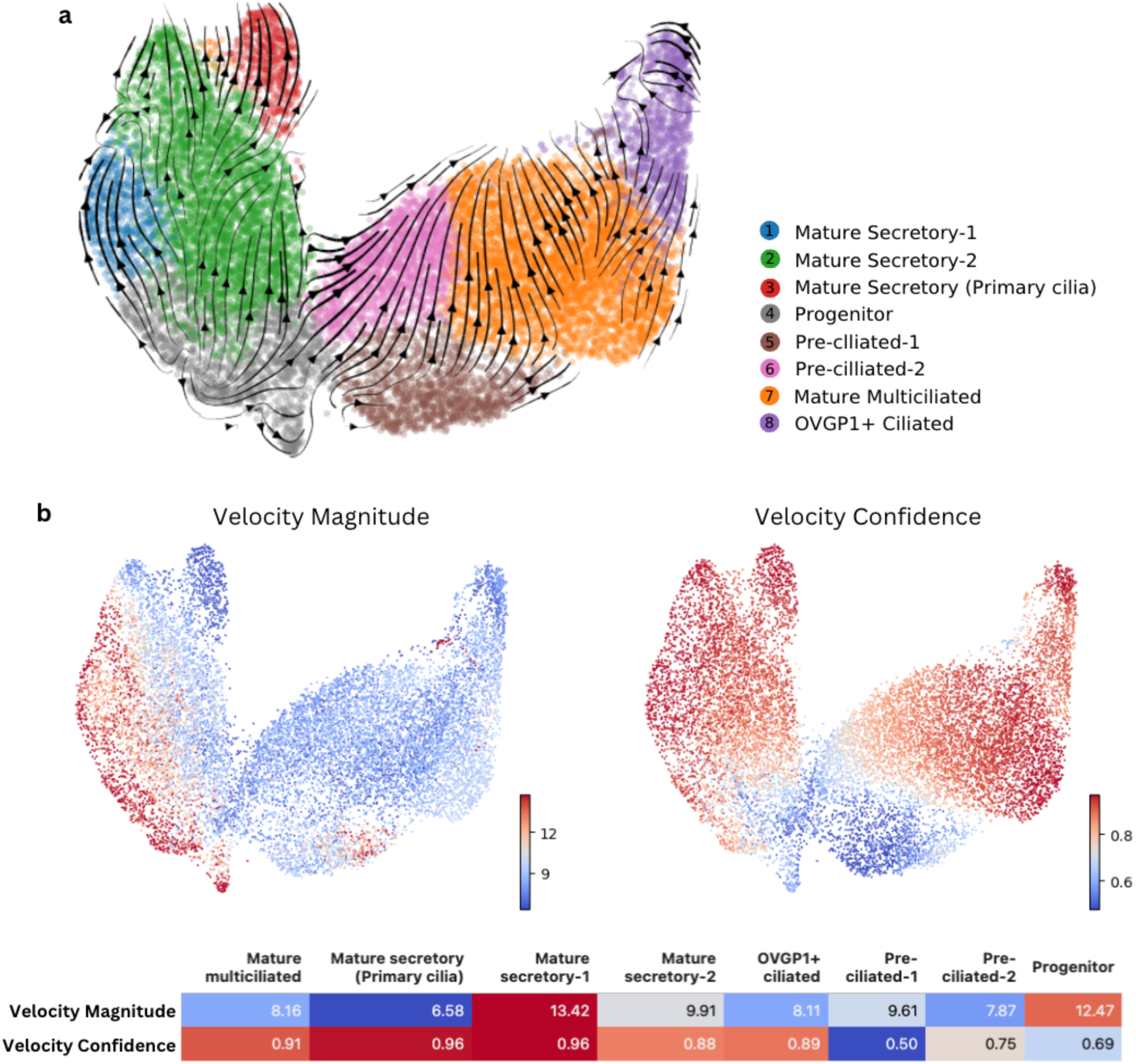
RNA Velocity Analysis of Fallopian Tube (FT) Epithelial Cells. **a**, Estimated RNA Velocity graph projected onto UMAP of FT epithelial cell populations, velocity vectors are visualised as streamlines. b, UMAP projections coloured by Velocity Magnitude, a measure of differentiation speed, and Velocity Confidence, a measure of agreement between the velocity vector of an individual cell and its neighbouring cell, for each cell, both values calculated per cell using scVelo. c, Table of average velocity length and velocity confidence for each FT epithelial cell population.

Velocity magnitude, a measure of differentiation speed, and velocity confidence, a measure of coherence between the velocity vectors of an individual cell and its neighboring cells, were also computed. (Fig3b). Higher velocity confidence and lower velocity magnitude are seen in Mature Ciliated and Mature Secretory populations, suggesting a cohesive differentiation trajectory at a slower pace, as expected for differentiated cell types [34, 35]. Developing Pre-Ciliated populations have a lower velocity confidence and a lower velocity magnitude, whilst the Progenitor population had the lowest velocity confidence and highest velocity magnitude for one population, suggesting chaotic and less coherent gene expression dynamics, consistent with that observed in other progenitor cell niches in vivo [34, 35].

### A Comparison of Fallopian Tube and Endometrial Epithelial Cells Reveals Transcriptome Differences Between Tissues

We next sought to identify the features that demarcate the FT from the endometrium, the developmental sister tissue and nearest anatomical neighbour. 58,463 endometrial epithelial cells were isolated from the raw endometrial scRNA-seq dataset, and epithelial populations annotated based on Garcia-Alonso et al. original data analysis[25] (Fig 4a).

**Figure 4.**
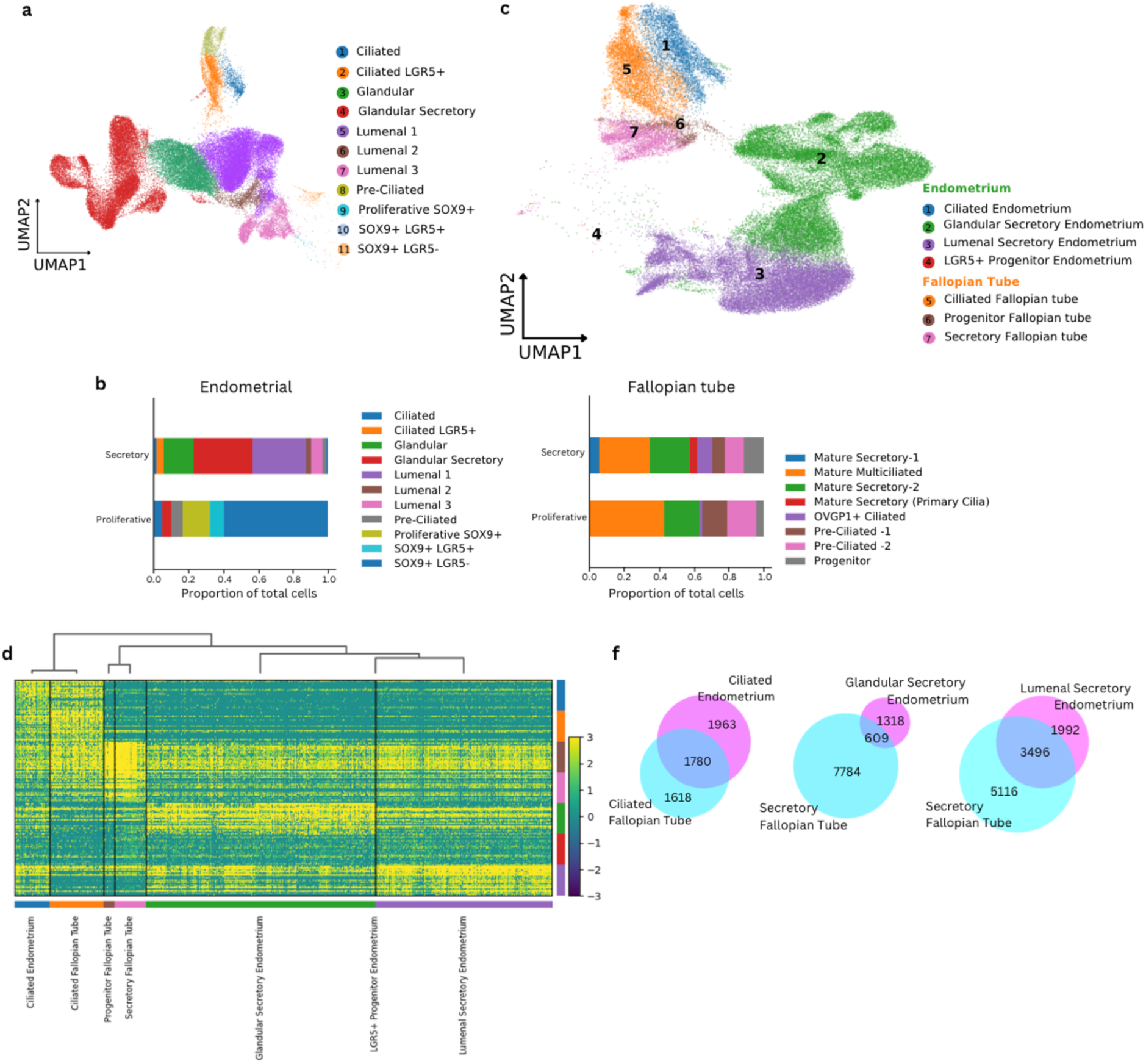
A Comparison of 57,951 Secretory Phase Fallopian Tube (FT) and Endometrial Epithelial Cells. **a,** UMAP of Epithelial cell populations isolated from Garcia-Alonso et al. Endometrial scRNA-seq dataset using scanpy workflow. b, Proportions of epithelial cell populations in the secretory and proliferative phases in the Endometrial scRNA-seq dataset, and the FT scRNA-seq dataset. c, UMAP projection of integrated endometrial and FT secretory phase epithelium, coloured by epithelial populations. d, Heatmap of top 25 differentially expressed genes (DEGs) for each epithelial population, genes presented on x-axis, and clusters of epithelial cell types on y-axis. f, Venn diagrams of significant DEGs (p<0.05) in ciliated and secretory epithelial cells of FT and endometrium.

Differences in the epithelial populations between secretory and proliferative phases were assessed between the FT and endometrium (Fig4b). Proportions of epithelial populations are similar between the proliferative and secretory phases in the FT (R2=0.800, P=0.003), unlike the endometrium where epithelial cell proportions change dramatically across the menstrual phase (R2=0.096, P=0.354) (FigS1). In the FT epithelium, Mature Secretory cells and *OVGP1+* Ciliated cells emerge in the secretory phase, suggesting a hormonal influence on differentiation.

Given these differences between menstrual phase, we next isolated and integrated secretory phase FT and endometrial epithelial cells, to develop a UMAP of 57,951 cells from 16 individuals. Clustering the integrated dataset revealed epithelial populations were well preserved (FigS2). To compare epithelial cells between the tissues, epithelial populations were grouped into 7 key categories, (1) Ciliated Endometrium; (2) Glandular Secretory Endometrium; (3) Lumenal Secretory Endometrium; (4) LGR5+ Progenitor Endometrium; (5) Ciliated Fallopian Tube; (6) Progenitor Fallopian Tube; (7) Secretory Fallopian Tube. FT and endometrial ciliated epithelial cells cluster together in UMAP projections and dendrogram clustering, suggesting similarity in global gene expression. Secretory epithelial cells remain distinct between the two tissues (Fig4c,d), suggesting transcriptomic differences between the secretory epithelial types of the endometrium and the FT.

Comparing significant differential gene expression (P=<0.05) between epithelial cell categories revealed that 7784 and 5116 genes are significantly differentially expressed in the FT secretory cells compared to endometrial glandular secretory cells and luminal secretory cells, respectively. In contrast, only 1618 genes were differentially expressed in ciliated FT cells compared to ciliated endometrial cells (Fig4f) (Supplementary Table 2).

### Functional Similarities and Differences Between Epithelial Secretory cells in Fallopian tube and Endometrium

Divergent gene expression between FT and endometrial secretory cells is suggestive of functional differences. Higher order network structures can identify genes causally linked to biological function. Here, hypergraphs were constructed from secretory menstrual phase expression correlation matrices. The top 600 differentially expressed genes between secretory phase FT and endometrial secretory cells were used as node genes, and the rest of transcriptome genes as edges in secretory epithelial cells of FT, glandular endometrium and luminal endometrium. Hierarchical clustering identified groups of 188 highly connected genes in the FT hypergraph, 108 genes in the glandular hypergraph, and 118 genes in the luminal epithelium hypergraph (Fig 5a,b,c).

**Figure 5.**
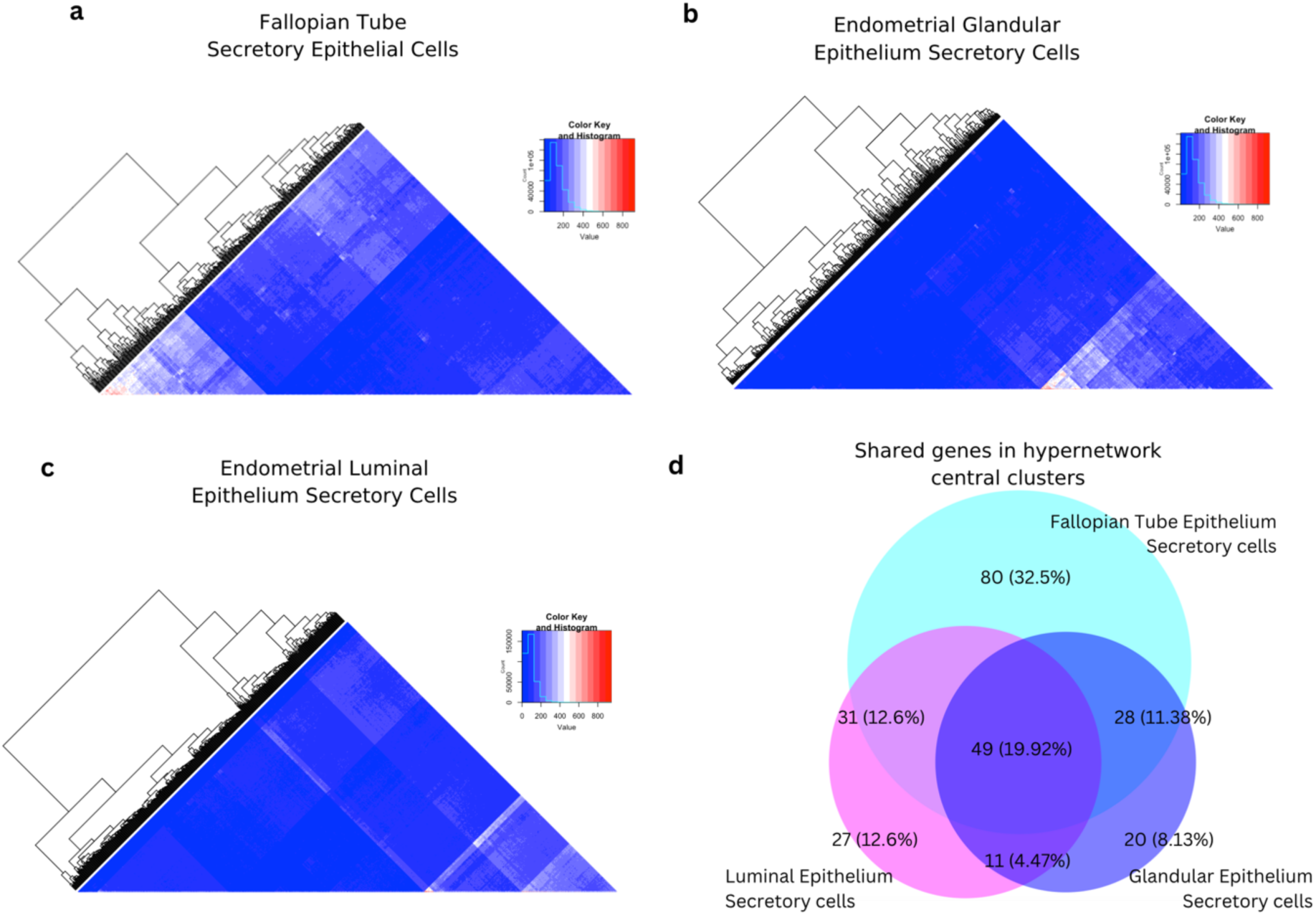
Hypergraph Analysis Identifies Genes with Functional Differences Between Secretory Cells in Fallopian Tube (FT) and Endometrial epithelia. Hypergraphs constructed using top 600 differentially expressed genes between FT and endometrial epithelial secretory cells, and edges as rest of transcriptome genes for secretory epithelial cells in a, FT. b, Glandular Endometrium. c, Luminal Endometrium. Higher values indicate a greater number of interactions between each differentially expressed gene and the rest of the transcriptome genes. d, Venn diagrams indicating number and percentage of unique and shared genes between hypergraph central clusters in each tissue.

Initial gene ontology over representation analysis of these clusters suggested functional DEG networks associated with lipid metabolism and immune response biological processes distinguish FT and endometrial secretory cells (FigS3.). Comparison between the tissue-specific hypergraphs revealed almost half (48.4%) of the clustered genes were shared, with a large FT-specific gene set (32.5%) compared to the luminal and glandular endometrium-specific subsets (11.0% and 8.1%, respectively) (Supplementary Table 3).

Examining the FT-specific gene set identified an over representation of glycoproteins *OVGP1, LCN2, SRGN, LRG1*, genes encoding cholesterol and lipid transport associated proteins, *APOA1, STARD4, STX11, SYNRG, TTC39B; and* several glycoprotein human leukocyte antigen class II genes, *HLA-DRB1, HLA-DRB5, HLA-DPA1 and HLA-DRA*.

### Hypergraphs Identify Tubal Ectopic Pregnancy Associated Genome Wide Association Genes

To establish whether functional networks predicted in the hypergraphs could be clinically relevant to FT pathology, we evaluated tubal ectopic pregnancy-associated genes from a recent GWAS [33]. From the lowest 10,000 p-value SNPs of the tubal ectopic pregnancy GWAS, 1338 rsIDs mapped to expression quantitative trait loci (eQTLs) in 22 of the 600 DEGs used to define our hypergraphs (Fig S3). Notably, 13 of these 22 genes clustered as the highest connected genes in the hypergraphs (Supplementary Tables 4,5). To quantify the importance of these 13 tubal ectopic pregnancy GWAS genes, hypergraph row sum, or degree centrality was calculated for each gene. Degree centrality is a measure of connectivity in a network, which represents the total number of interactions between the DEGs gene and other genes [36].

Several ectopic pregnancy related GWAS genes were highly connected to the rest of the transcriptomes of all three tissues, *SLC7A2, MUC1, CLDN1 and GLS* (Fig 6a). MUC1 was the strongest genetic association previously identified, thus validating high connectivity in the hypergraph as a clinically relevant functional metric. These genes are indicative of shared functionality between tissues. *HLA-DRB1* and *HLA-DRB5* are ectopic pregnancy GWAS genes that were highly connected only in the FT secretory cell transcriptome. Additionally, *ANKRD36B. CLGN, PEX6, PLXNA4* and *NR2F1* were highly connected only in the luminal endometrial secretory cells while *ANKRD36* and *PGGHG* were highly connected in the glandular endometrial secretory cells. *MUC1* and *SLC7A2* also had the highest expression in Mature Secretory-2 FT cells (Fig6b).

**Figure 6.**
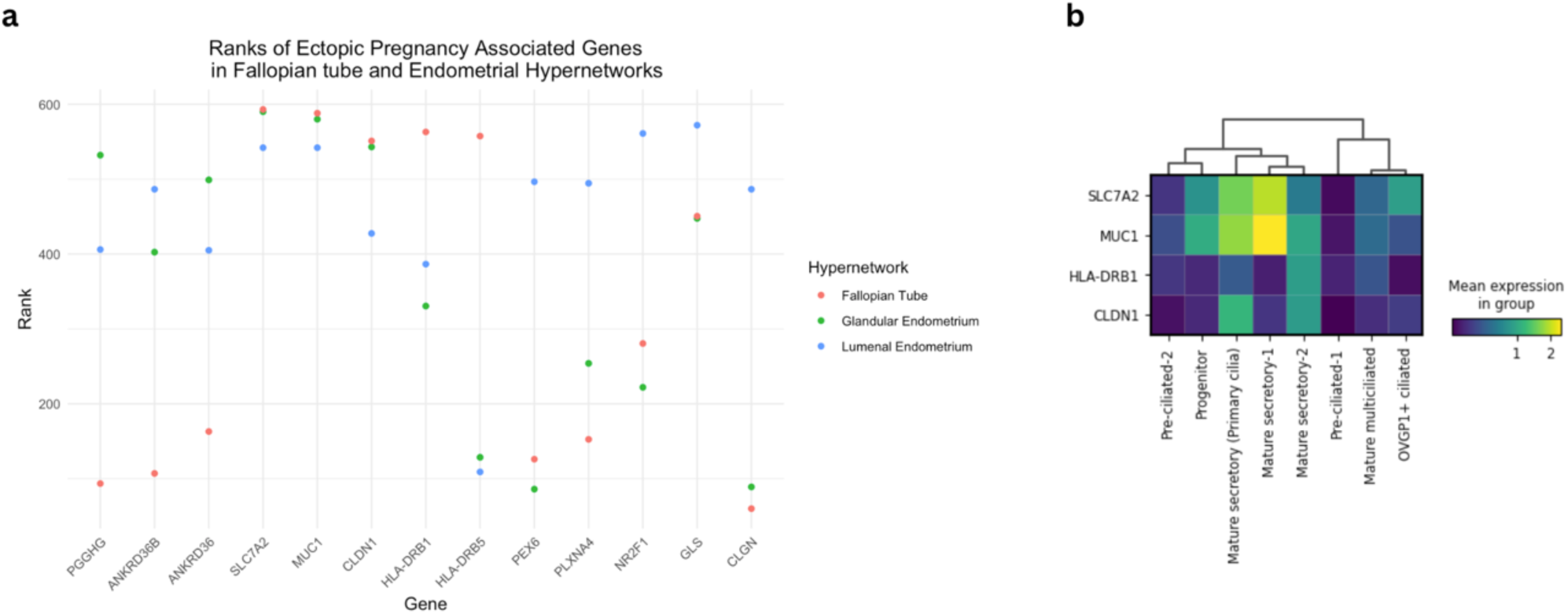
Tubal Ectopic Pregnancy Associated Genome Wide Association Study (GWAS) Genes are Ranked Differentially in Hypergraphs of Secretory Epithelial Cells of the Fallopian Tube (FT), Glandular and Luminal Endometrium. a, Ranked rowsums of tubal ectopic pregnancy genes in secretory epithelial cell hypergraphs for FT, glandular endometrium and luminal endometrium. b, Mean Expression of 4 highest ranked GWAS genes in FT epithelium.

## Discussion

The epithelial lining of the FT regulates key functions of the organ; gamete and embryo transport, and pre-implantation embryo development support. Our meta-analysis of published FT scRNA-seq data characterised 8 FT epithelial populations, and mapped a previously unseen bi-directional differentiation of ciliated cells and specialised mature secretory cells from a *OVGP1+* progenitor population. This differentiation to mature cell types coincides with the secretory phase of the menstrual cycle. Differences between FT epithelium and that in the neighbouring endometrium were predominantly in the secretory cell populations, with hypergraph analysis identifying lipid metabolism and immune interaction pathways as functional divergences in FT secretory cells. The functional importance of hypergraph genes was underscored by their prominence in tubal ectopic pregnancy GWAS, reasserting the causal involvement of MUC1 in implantation, and suggesting new genes that contribute to normal and pathological implantation.

By performing a meta-analysis of healthy FT epithelial cells we identify 8 secretory, ciliated and progenitor cell types, and map their bi-directional differentiation trajectory. Here the progenitor population is characterised by *OVGP1* and *CRISP3* expression, and absence of both mature secretory (*CXCL2, C3, PAX8*) and ciliated cell markers (*FOXJ1, PIFO*). Earlier studies have reported linear differentiation trajectories of FT epithelium [18, 19]. Dinh et al. construct a linear trajectory using pseudotime analysis, where secretory clusters differentiate to ciliated cells via a population of ‘uncharacterised cells’, which closely resemble our populations of Pre-Ciliated-1 and Pre-Ciliated-2.

Conversely, Ulrich et al. describe two epithelial progenitor populations where a ‘Peg Cell’ differentiates to secretory cells, and an ‘EMT progenitor’ gives rise to ciliated cells. Significantly increasing the number of cells analysed and utilising RNA velocity analysis allowed us to map a bi-directional differentiation model where the Progenitor population forms either Mature Secretory or Mature Multiciliated cells. This differentiation coincides with the menstrual phases, where the mature populations are only present in the secretory phase. Previous studies have demonstrated the FT does not undergo cyclical proliferation but it is hormonally responsive [37]; these results challenge this assumption and suggest it may indeed undergo cyclical maturation and differentiation.

We validated our development model using RNA velocity confidence and magnitude which previous FT differentiation trajectory studies have not done. Based on the principles of gene expression dynamics, chaotic gene expression suggests a progenitor or pluripotent stem cell type[34]. In the ‘Progenitor’ population, high velocity magnitude indicates a faster differentiation trajectory, and low velocity confidence indicates velocity vectors are not correlated between neighbouring cells and are therefore more chaotic, suggesting a pluripotent cell type [29, 35]. Development of stability, and loss of chaos in gene expression is also associated with loss of pluripotency, and terminal differentiation [38]. This is seen in the mature epithelial populations where velocity has a smaller magnitude and higher confidence compared to the progenitor population. This suggests differentiation trajectories have slowed, and there is a cohesive agreement between neighbouring cells in the direction of velocity vectors, or development.

In vitro studies have supported this FT epithelial cell differentiation model where a secretory-like progenitor cell, gives rise to ciliated and mature secretory cells. Yamamoto et al. 2015, characterised FT epithelial cells from primary human adult and fetal FT cells using clonal isolation techniques.

These stem cells did not express *LGR5*, or secretory or ciliated markers *FOXJ1* or *PAX2*, and underwent differentiation in a trans-well air liquid interface culture to form secretory and ciliated cells [39]. Suggesting a pluripotent population of *LGR5-* FT epithelial cells exists, similar to our ‘Progenitor’ cells here. Assessing the progenitor population in vitro is critical to confirming this model of FT epithelial differentiation.

We compared secretory phase FT and endometrial epithelial cells to determine how they contribute to tissue-specific function. Previous studies have compared expression of individual transcripts and proteins between the two tissues [37, 40–43], but there is no comparison of transcriptomes at a bulk or single cell resolution. As observed across human tissues, ciliated cells in the FT and endometrium have a highly similar transcriptomes [44].

In contrast, large differences were seen between FT and endometrium secretory epithelial cells. By constructing hypergraphs to identify higher-order gene interactions[20, 21] we identified clusters of genes likely to underpin functional tissue divergence. The widely used FT-specific marker OVGP1 was among these genes, suggesting that this secreted glycoprotein has functional roles in FT that have previously been ascribed to interactions with the zona pellucida that regulate fertilisation and embryo development [45, 46]. LCN2, another secreted protein specific to the FT hypergraph, is expressed throughout the female reproductive tract and is canonically involved in iron transport.

However recent studies have suggested roles for LCN2 in regulating the innate immune system in the endometrium [47]. Lipid and cholesterol transport associated genes were also identified as functional genes, *APOA1, STARD4, STX11, SYNRG, TTC39B.* Lipids are known to constitute a component of FT fluid and thought to influence oocyte maturation and embryo development [48, 49]. Several of these genes are also associated with intra and extracellular vesicle transport which has been described as a component of FT fluid [50–53]. These functional genes describe a secretory, lipid rich environment facilitated by the highly specialised secretory cells of the FT.

As the hypergraph models identified genes previously shown to have functional roles in the FT, we wanted to assess whether genes relevant to tubal ectopic pregnancy, a common FT pathology, were also identified. Remarkably, of twenty-two tubal ectopic pregnancy GWAS genes present in the hypergraph DEGs, thirteen were found to be in the most connected gene clusters including the highest scoring disease-associated gene, MUC1. Ranking these GWAS genes based on their row sum, or connectivity, in the hypergraph models offered a metric for functional importance and yielded other genes likely to be causally involved in tubal ectopic pregnancy.

Notably, *MUC1* was the second highest ranked GWAS gene in the hypergraphs, providing confidence that this metric was correlated to clinical and functional importance. Furthermore, this analysis may shed light on eutopic implantation as other genes suspected of mechanistically supporting embryo attachment and implantation were present in the ectopic pregnancy GWAS and hypergraph central clusters of the FT and luminal endometrial epithelium; *CLDN1* and *PLXNA4* [54–56]. The strong association of tubal ectopic pregnancy and implantation in the secretory cells of FT and endometrial luminal epithelium also suggests that specialised secretory cells mediate embryo attachment in health and disease. *HLA-DBR1* and *HLA-DRB5* were highly connected in the FT. HLA alleles have also been associated with recurrent implantation failure and miscarriage [57, 58].

When we examined expression levels of the ectopic GWAS genes, *SLC7A2* and *MUC1* are both expressed highest in the Mature Secretory-1 FT population. Interestingly, this mature epithelial population is only present in the secretory phase. These specialised secretory cells may protect against ectopic embryo implantation in the secretory phase and ensure the embryo is transitioned to the embryo by providing a unique secretory environment with high levels of MUC1.

In summary this work has demonstrated the utility of sophisticated meta-analyses of existing data sets to reveal novel insights into the tissue structure and dynamics of the FT. As well as yielding novel insights into the differentiation process of mature epithelial cell types, our approach has identified candidate genes of clinical relevance in the susceptibility of individuals to ectopic implantation.

## Supplementary figures

**Supplementary Figure 1.**
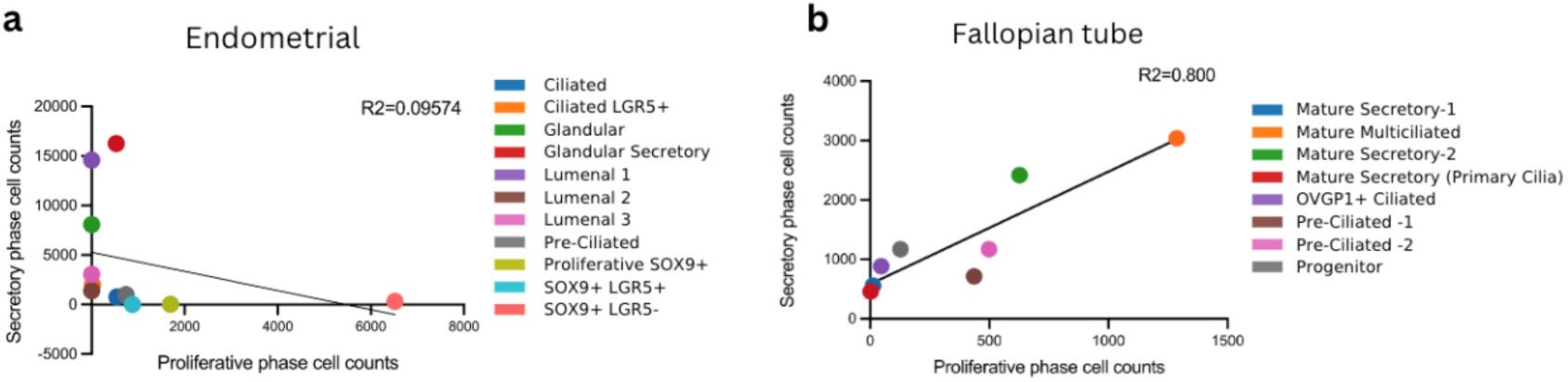
Endometrial and Fallopian Tube (FT) Epithelial Cell Population Counts in the Secretory Versus Proliferative Menstrual Phases. **a**, Linear regression of endometrial cell counts between menstrual phases, R2=0.09574. b, Linear regression of FT cell counts between menstrual phases, R2=0.800.

**Supplementary Figure 2.**
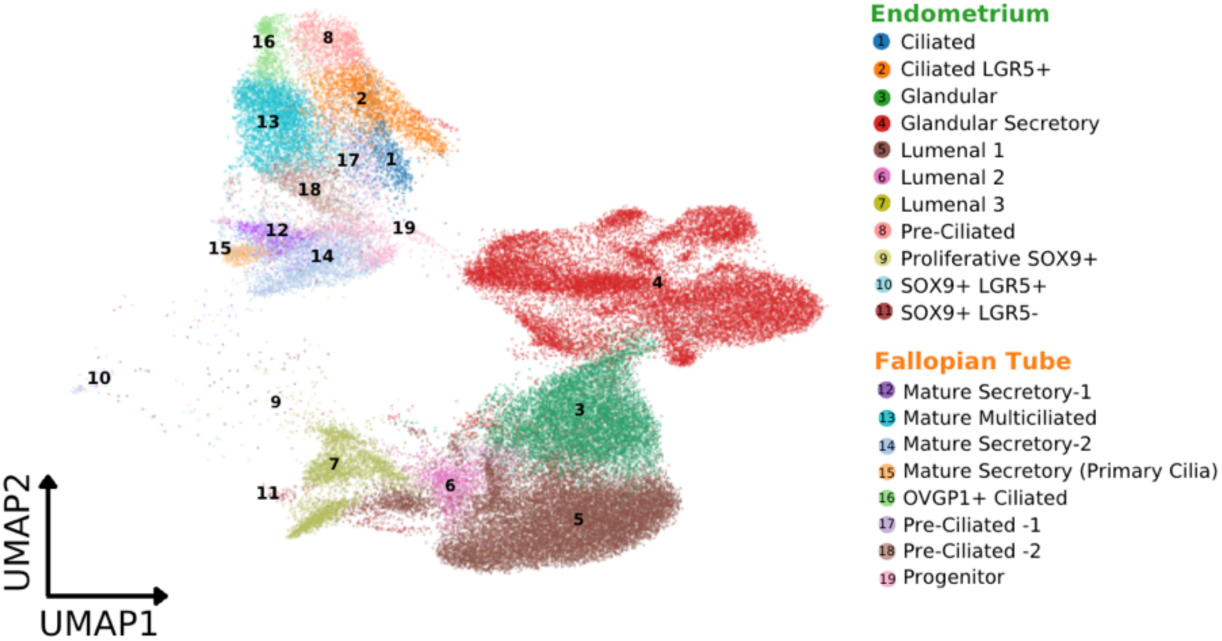
UMAP of Fallopian Tube and Endometrial Epithelial Cells in the Menstrual Secretory Phase. All epithelial populations are shown.

**Supplementary Figure 3.**
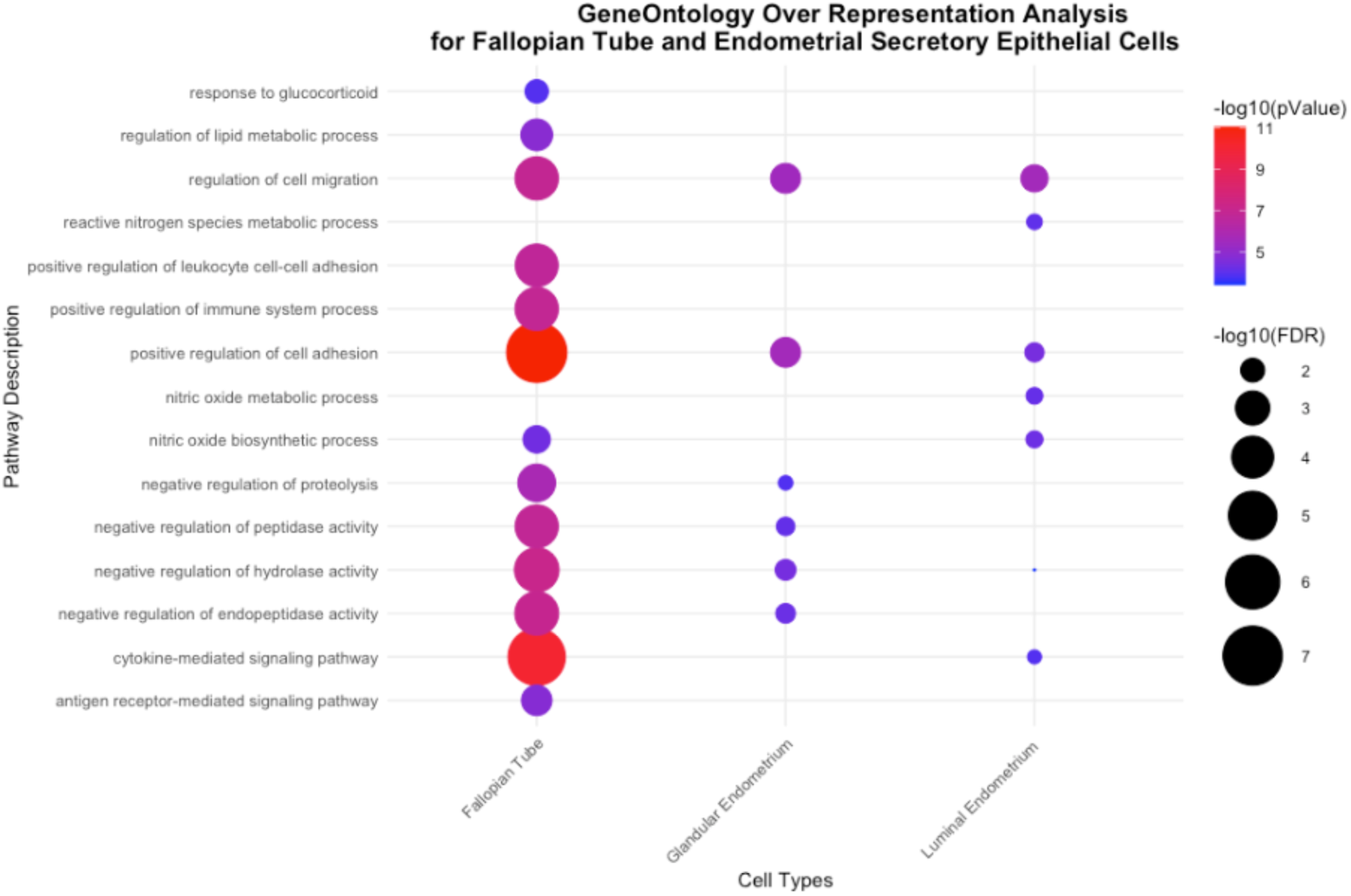
Dot plot of GeneOntology Over Representation Analysis using Central Clusters of Fallopian Tube, Glandular and Luminal Endometrium Hypergraphs. False discovery ratio (FDR) and pValue are shown as -log10.

**Supplementary Figure 4.**
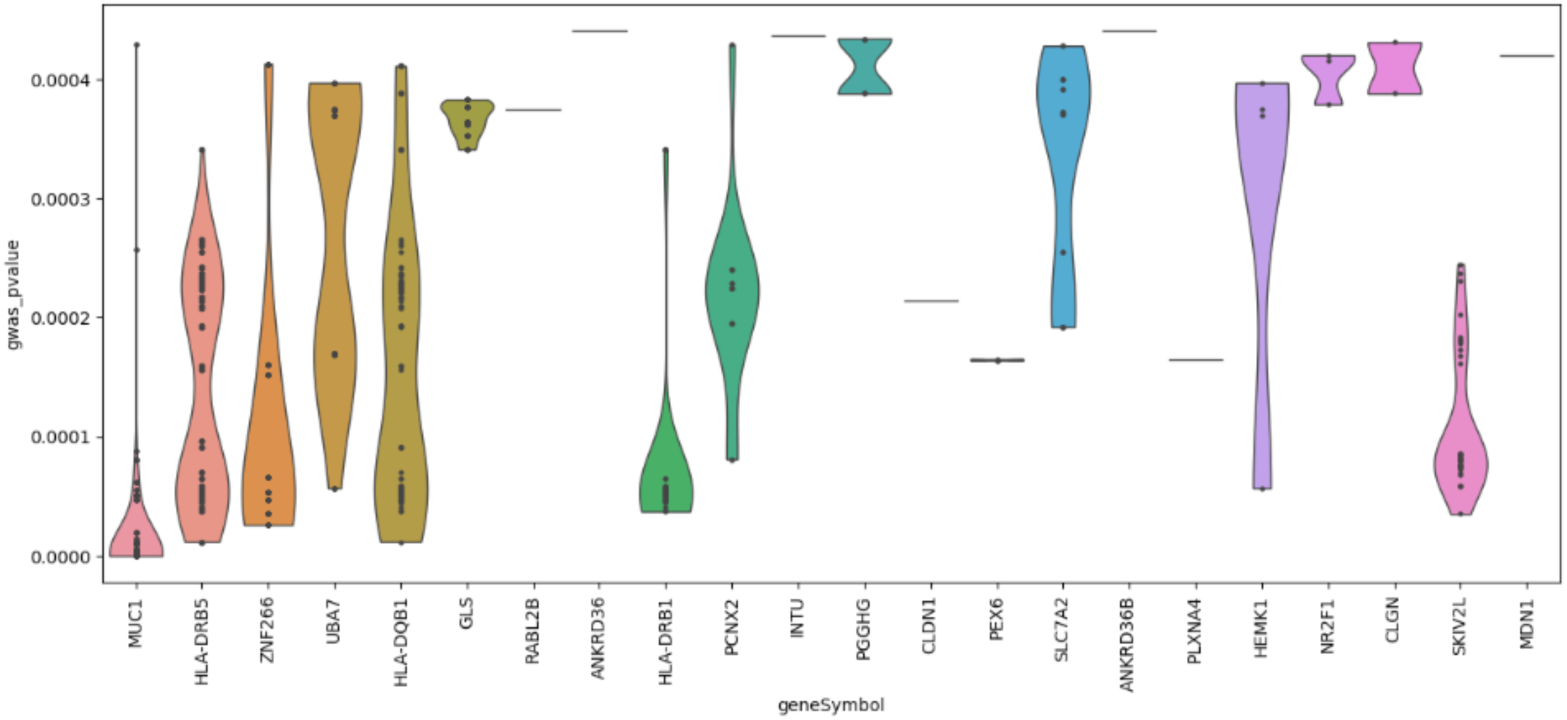
Violin Plot of the Ectopic Pregnancy Genome Wide Association Study (GWAS) Genes pValues. pValues (P = 1.68 x 10^-21^– 5.8 x 10^-4^) of GWAS genes in hypergraph models, from the summary statistics in gwas catalog, study accession GCST90272883, each point represents pValue of individual rsID.

## Notes

### Competing Interest Statement

The authors have declared no competing interest.

